# Integrated Single-Cell and Spatial Profiling of Matrix Metalloprotease Gene Expression in Colorectal Cancer

**DOI:** 10.64898/2026.04.17.719089

**Authors:** Nicholas A. Danese, Shan Kurkcu, Marina Bleiler, Klea Nito, Alan Kuo, Daniel W. Rosenberg, Masako Nakanishi, Charles Giardina

## Abstract

Increased matrix metalloproteinase (MMP) expression has long been recognized as a common feature of colorectal cancers (CRCs), yet less is known about how these enzymes interact to impact cancer progression. Of particular interest is MMP expression overlap where multi-substrate targeting can accelerate extracellular matrix (ECM) breakdown. Taking advantage of single-cell and spatial transcriptomic data, we analyzed MMP expression in CRCs, focusing on cancer-associated fibroblasts (CAFs), myeloid cells, and cancer cells. Single-cell RNA-seq data revealed that distinct CAF subtypes express different MMP combinations, including MMP1/2/3 and MMP2/11-expressing CAFs. Evidence was also obtained that the number of MMP1-expressing CAFs in CRCs is highly variable, likely impacting stromal ECM dynamics. In contrast to CAFs, myeloid cells (monocytes, macrophages, and dendritic cells) all expressed varying levels of the same "myeloid MMPs" 9, 12, and 14, with MMP expression correlated with a cell’s secretory potential, a phenotype that would facilitate MMP release. Finally, a subpopulation of cancer cells expressed high levels of MMP7, which was frequently co-expressed with several Wnt-related genes. Analysis of spatial transcriptomic data showed MMP expression clusters, driven largely by cell-type, with regions of notable MMP expression overlap. High resolution spatial analysis revealed clusters of cells at the cancer-stroma interface expressing a broad spectrum of MMPs. These multi-MMP expression clusters included cancer cells, CAFs, and myeloid cells all expressing MMP7 and a broad range of other MMPs. These multi-MMP foci also exhibited elevated Wnt signaling and EMT markers and may represent sites from which cancer cells escape the primary lesion.

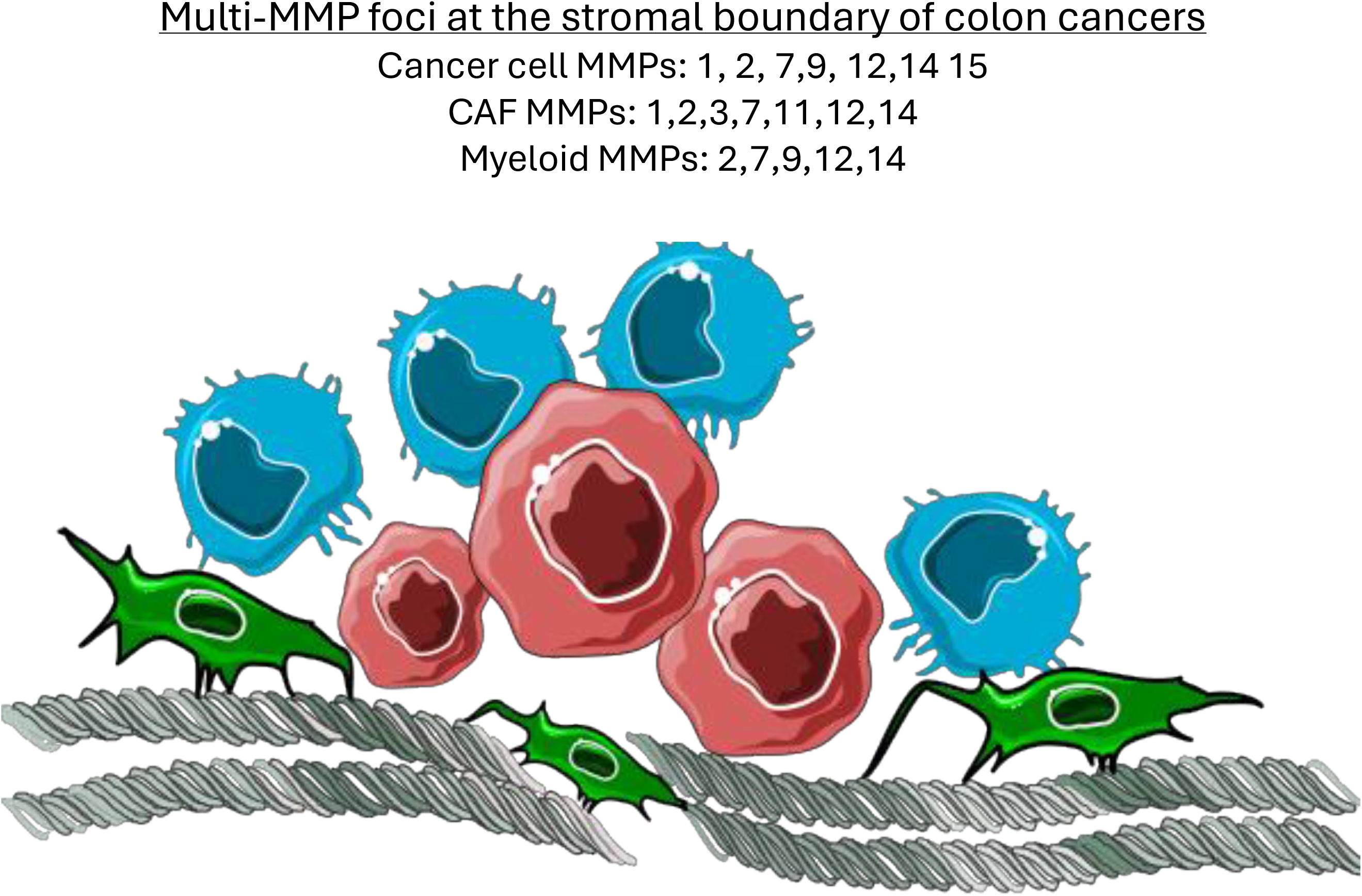

## Introduction

Colorectal cancer (CRC) is the third most diagnosed malignancy worldwide and a leading cause of cancer-related mortality, with an estimated 1.9 million new cases and over 900,000 deaths annually [1]. While the overall incidence of CRC is lowering, it is increasing in younger populations, particularly in high-income countries, emphasizing the need for improved understanding of CRC etiology and pathophysiology [1, 2]. CRCs consist of both genetically transformed epithelial cells and a diverse stromal compartment comprised of fibroblasts, endothelial cells, and inflammatory cell infiltrates. Cancer cells exhibit uncontrolled proliferation and invasive capacity, while stromal cells serve as key modulators of tumor progression in part through their roles in angiogenesis and immune and inflammatory responses [3]. Reciprocal interactions between cancer cells and stromal components actively shape the tumor microenvironment (TME), which can significantly impact cancer growth and progression.

Matrix metalloproteinases (MMPs) are a family of zinc-dependent endopeptidases that mediate extracellular matrix degradation and thus play a pivotal role in tissue remodeling under both physiological and pathological conditions. MMPs cleave ECM components, which in turn can affect cell attachment, signaling, and movement through a tissue. MMPs can therefore affect myriad events in cancer, ranging from metastasis to immune surveillance. These enzymes are classified based on their substrate specificity and structural characteristics into collagenases (MMP1, MMP8, MMP13, MMP18), gelatinases (MMP2, MMP9), stromelysins (MMP3, MMP10, MMP11, MMP12), matrilysins (MMP7, MMP26), and membrane-type MMPs (MT-MMPs, e.g., MMP14, MMP15, MMP16, MMP25)[4, 5]. Each group cleaves a selection of ECM components, with some overlap between groups. MMP expression increases in inflamed and neoplastic tissues, with much of this regulation occurring at the transcriptional le vel. MMPs are initially expressed as zymogens (pro-MMPs) that require cleavage for activation, often by other MMPs, and are inhibited by tissue inhibitors of metalloproteinases (TIMPs), providing two additional layers of regulation. MMP cleavage by other MMPs establishes activation cascades, wherein one MMP activates another. For example, MMP3 can activate both MMP7 and MMP9, while MMP2 is activated by MMP14 [6–8]. MMP7 plays a key role in CRC by degrading basement membrane components and activating MMP2 and MMP9, reinforcing a proteolytic network that sustains tumor progression [6]. This cooperative enzymatic interplay between MMPs enhances ECM degradation to affect cellular trafficking and metastasis. Understanding the interplay between various MMP subtypes within CRC tissue is therefore important for understanding numerous events of CRC progression.

While MMPs have been extensively studied, their spatial expression within CRC (and other cancers) remains less well defined. Advances in single-cell RNA sequencing and spatial transcriptomics now enable high-resolution mapping of MMPs in CRC, providing insights into their localization and potential functional interactions. This knowledge could lead to a better understanding of how these enzymes collectively influence cancer progression. This information could also be leveraged to develop novel therapeutic approaches that exploit MMP expression either through inhibitors that halt lesion growth or as targets for cancer-seeking therapies and prodrugs.

## Materials and Methods

### Data acquisition and processing

Bulk RNA-seq data for identifying MMPs were obtained from the TCGACOAD STAR cohort. Single-cell RNA-seq data were retrieved from GEO (GSE132465), and spatial transcriptomic datasets were obtained from 10X Genomics (CytAssist_11mm_FFPE_Human_Colorectal_Cancer, Tumor 1; Visium_HD_Human_Colon_Cancer, Tumor 2). All analyses were performed in R. Bulk RNA-seq counts from the TCGACOAD cohort were normalized using DESeq2, and differential MMP expression between cancer and normal tissues was visualized with ggplot2. Highly expressed MMPs were further examined in single-cell and spatial datasets. Single-cell data were processed as Seurat objects using SCTransform, with celltype annotation via SingleR using the HumanPrimaryCellAtlas dataset (restricted to CRC-relevant types) as reference. Data were subset into epithelial, myeloid, and fibroblast/smooth-muscle compartments. Clustering followed guidelines established by Satija Lab’s Guided Clustering Tutorial, using Seurat’s RunPCA, FindNeighbors, and FindClusters functions. Clustering was carried out based both on variable genes and selected MMP genes. Nearest-neighbor and shared-nearest-neighbor approaches across multiple resolutions provided complementary cluster perspectives.

### Spatial data processing

The spatial transcriptomic data from the two colon cancers were captured at different resolutions. To make the datasets comparable in pixel number and reads per pixel, the 8 gm • 8 gm bins of tumor 2 were aggregated into 56 gm • 56 gm bins. Both tumors were processed as Seurat objects and clustered using variable gene and MMP-based approaches. Pixel level cell-type assignments used SingleR with the same reference as single-cell data. Fibroblast and smooth-muscle–associated pixels were further subset for focused analyses.

### Visualization and downstream analyses

Dot plots, heatmaps, violin plots, and bar charts were generated with ggplot2. Cells were classified as positive or negative for a gene based on the median expression of all expressing cells. SpatialFeaturePlot and SpatialDimPlot were used to visualize spatial MMP expression and clustering. One additional gene expression heatmap was created with ComplexHeatmap. Cluster-defining genes were identified using Seurat’s FindAllMarkers function. The top 500 positive markers from each cluster were analyzed with PANTHER Gene Ontology for Biological Process. Enrichment, probability, and representative processes were summarized in gt-generated tables.

### Fibroblast isolation and culture

Colorectal cancer tissue specimens were obtained from UConn Health/Waterbury Hospital. All samples and associated clinical data were collected from consented patients undergoing colon resection and were de-identified in accordance with institutional protocols and established ethical guidelines. Cells were tested for presence of mycoplasma and their absence was confirmed. Tissues were washed with 1x Hanks’ Balanced Salt Solution and incubated with 25 mM ethylenediaminetetraacetic acid (EDTA) for 1 hour at 4o C. Epithelial cell layer was stripped off by multiple rounds of vortexing. Remaining tissues were incubated with collagenase solution (1 mg/ml collagenase, 1 mg/ml dispase and 10ug/ml DNase I) for 90 min at 37o C. Digested tissues were filtered (70 mm), centrifuged at 300g for 5 min and washed with Dulbecco’s Modified Eagle Medium (DMEM) media supplemented with 10% fetal bovine serum (FBS), 2 mM L-glutamine and 1x Penn/Strep solution (Gibco). Resuspended cells were cultured in complete DMEM, and frozen in aliquots with complete medium plus 10% dimethyl sulfoxide. For growth and analysis, frozen cell aliquots were thawed and maintained in Lonza fibroblast growth medium (FGM-2 Fibroblast Growth Medium-2 BulletKit) at 37°C in a humidified atmosphere with 5% CO2. Cells were subcultured for up to seven passages and monitored for the changes in morphology. Cells were checked for mycoplasma contamination and confirmed to be negative.

### Fibroblast treatments and analyses

To assess MMP1 expression in response to stimuli, cells were grown on coverslips, serum-starved overnight, and then treated with 10 ng/mL TNF (Thermo Fisher Scientific), 1 gg/mL LPS-EB (InvivoGen), or graded serum levels for 20 h. To assess differences in MMP1 expression between fibroblast colonies, cells were seeded on coverslips in 24-well plates at a density of 120 cells per well with the addition of conditioned medium collected from the cells’ previous plate (25% v/v). Cells were incubated for 9 days, with medium replenished as needed. For the RNA-seq analysis, RNA was isolated using RNeasy extraction kit (Qiagen), according to manufacturer’s instructions. RNA quality assessment was performed at the Center for Genome Innovation (CGI) core facility at the University of Connecticut. Library preparation and bulk RNA sequencing were performed by Azenta Life Sciences on an Illumina NovaSeq 6000 platform, generating 150 bp paired-end reads with a target depth of ∼30x. Data were normalized and analyzed using DESeq2 (R/Bioconductor) to quantify gene expression changes. For immunofluorescence staining experiments, cells were fixed in 3% paraformaldehyde, permeabilized with 0.3% Triton X-100 in the Dulbecco’s phosphate-buffered saline (DPBS), and blocked in 5% Goat serum (Jackson ImmunoResearch) in DPBS. Cells were then stained for MMP1 expression using the GeneTex MMP1 antibody GTX100534 and appropriate secondary antibodies (Thermo Fisher Scientific, Jackson ImmunoResearch). Nuclei were visualized using DNA dye DAPI (Thermo Fisher Scientific). Fluorescent images were acquired with a Nikon A1R laser scanning confocal microscope (Nikon Instruments, Japan) and analyzed using Fiji /ImageJ software. Two-sided Fisher’s exact test (GraphPad Prism) was used to compare MMP1–positive cells between normal fibroblasts and CAFs; p < 0.05 was considered significant. Data were presented as percentages. Secreted MMP1 protein levels were quantified via Western blot using Mini-PROTEAN TGX precast gels (Bio-Rad), GeneTex MMP1 primary antibody, and IRDye® 680RD secondary antibody. Blots were visualized on a LiCor Odyssey® CLx Imaging System.

## Results

### MMP expression in CRC

TCGA data was analyzed to identify the MMPs highly expressed in CRC. This analysis revealed high expression of MMPs 1, 2, 3, 7, 9,11,12,14 and 15 (Fig 1A). Most of these MMPs are expressed at higher levels in cancer than normal tissue, although MMPs 2, 14, and 15 are expressed at relatively high levels in normal tissue. Since CRCs are populated by numerous cell types that can impact lesion progression, we determined the MMP expression profiles in different cell types within healthy and cancerous tissues using scRNA-seq data and the HumanPrimaryCellAtlas for reference to determine cell type. In normal tissue, scRNA-seq showed fibroblasts (FBs) and smooth muscle cells (SMCs) to be the highest MMP expressors, with MMP2 being the highest expressed isoform (Fig 1B). In CRC, many more cell types express MMPs; there are notable increases in epithelial cells (with MMP7 predominating) and myeloid cells (with MMPs 9, 12, and 14 predominating), in addition to an increased expression in fibroblasts (Fig 1C). A table summarizing cell-type MMP expression in normal and CRC tissue is shown (Fig 1D).

**Figure 1:**
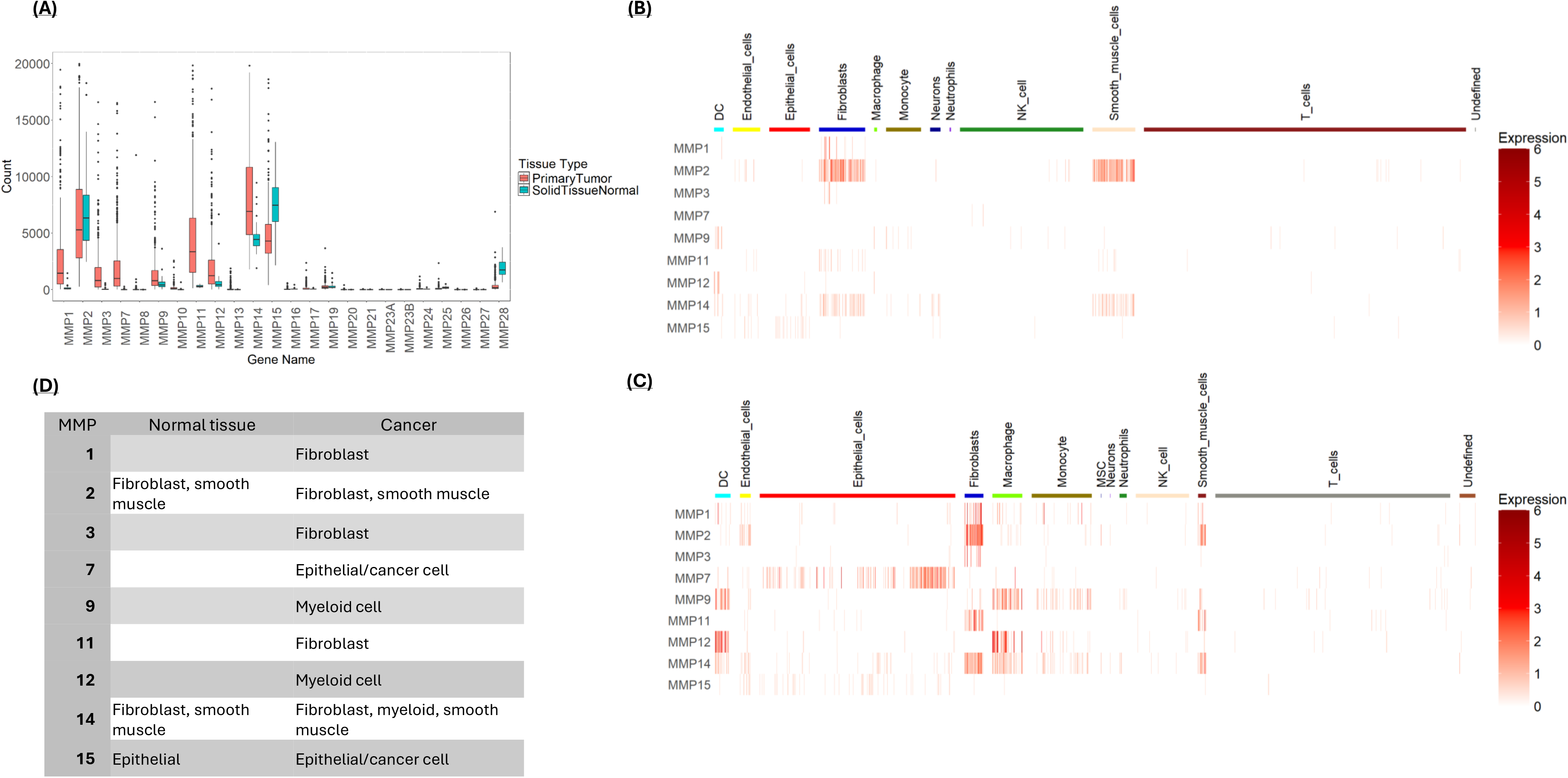
A) MMP expression in colorectal tissue. Data was retrieved from TCGA and analyzed to identify the most highly expressed MMPs in noncancerous mucosa and primary CRC lesions. Shown are the normalized counts from RNA-seq data. **B&C)** Single cell RNA-seq (scRNA-seq) data was used to determine MMP expression patterns in cell types isolated from normal colon tissue **(B)** and cancerous colon tissue **(C)**. Single cell RNA sequencing was obtained from the NCBI GEO database (accession code GSE132465). Normalized expression levels of MMPs are shown in rows with cell type indicated by the color bar across the top of the heatmap. **D)** Comparative cellular expression of MMPs in normal and tumor microenvironments summarizing the results presented in **1B** and **1C**.

### MMP expression in CAFs and SMCs

Since CAFs and SMCs are related cell types, and the CRC single-cell data showed them expressing a similar set of MMPs, these two cell types were combined and resolved into 8 clusters (Fig S1). The SMCs and CAFs equally populated many of the clusters, consistent with these two cell types expressing similar MMPs. However, clusters 3, 4 and 5 were mostly CAFs and were distinguished by high MMP1/3, high MMP11, and high MMP1 expression, respectively. To determine whether the MMP expression pattern was stochastic or associated with distinct cell subtypes, we analyzed differential gene expression within the MMP-defined clusters (Fig S2). This analysis showed that these clusters also differentially expressed other genes, consistent with a preferred MMP expression pattern for different CAF and SMC subtypes. Since multiple different CAF types have been described, “fibroblasts” were pulled out of the data set and analyzed separately. CAFs were resolved into three clusters based on their MMP expression, including an MMP1/3 cluster and an MMP11 cluster (Fig 2A). The top genes enriched in these clusters are shown in Fig 2B. Cluster 0 cells expressed numerous SMC markers and were identified as SMCs by ToppCell Atlas, suggesting a population of CAFs with SMC characteristics. The MMP1/3 CAFs showed high chemokine expression, especially neutrophil chemokines and were identified as “mesenchymal” fibroblasts by ToppCell Atlas (Fig 2B). Finally, MMP11-expressing CAFs showed expression of numerous secreted ECM components and were identified as “myofibroblasts”. This analysis is consistent with different CAF subtypes having preferred MMP expression profiles, which likely impacts their role in regulating ECM dynamics in the tumor [9].

**Figure 2:**
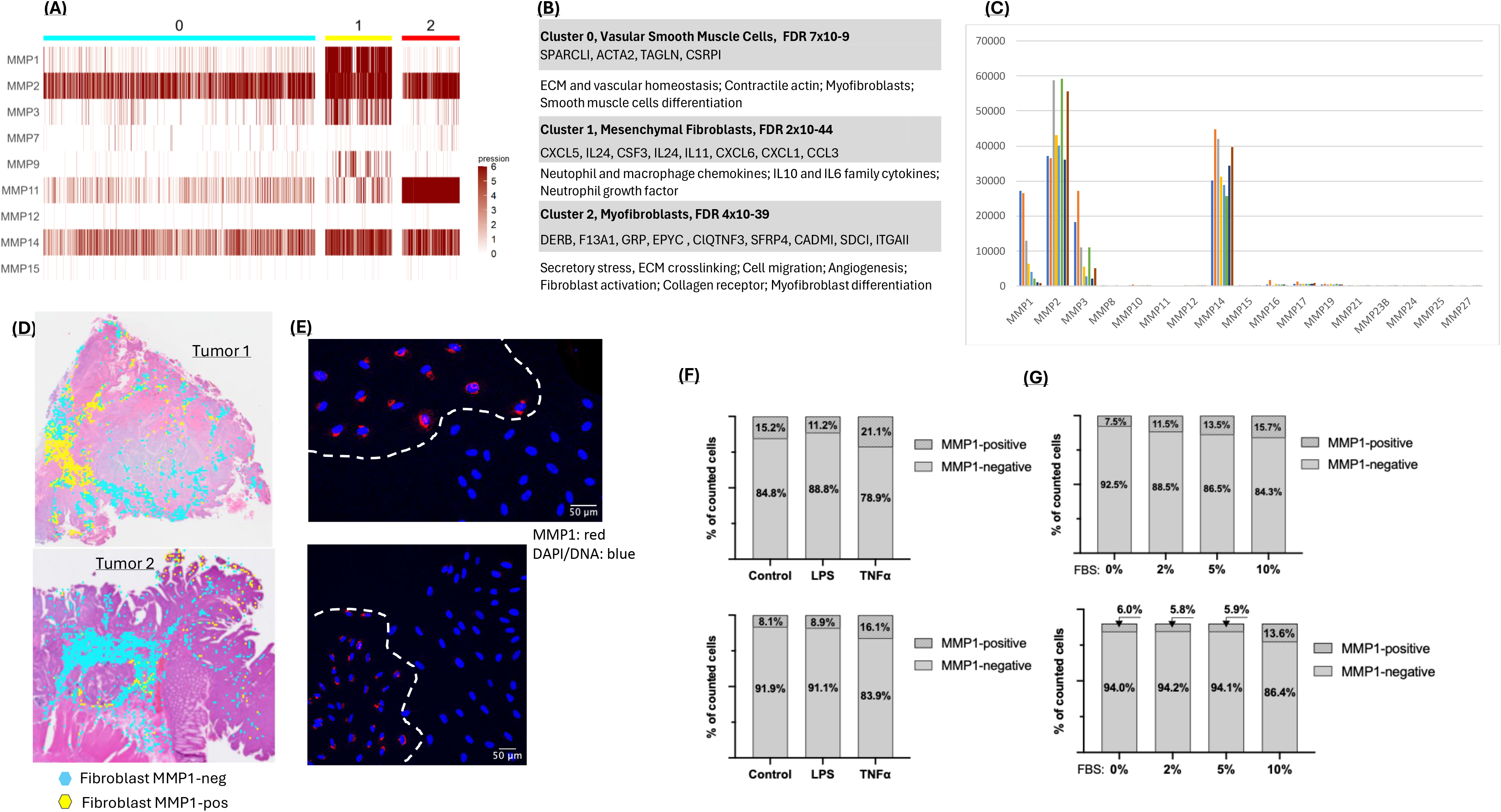
A) CAFs from CRC were clustered based on their MMP expression to a resolution that generated 3 groups. Expression of MMPs across these groups is displayed. **B)** Genes and CAF types correlating with MMP expression patterns. Genes that were >2-fold enriched in these MMP-defined CAF clusters are shown, together with the correlating cell type based on ToppCell Atlas analysis. **C)** MMP expression in eight primary CAF cell lines established from human colon cancers as determined by RNA-seq. **D)** CAFs were identified in spatial data from two CRCs. Pixels showing expression of CAF-related genes are shown, with MMP1-positive CAFs shown yellow and MMP-negative CAFs shown in cyan. **E)** CAFs plated at low density to form discrete colonies were stained for MMP1 expression. MMP1 staining is shown red and DAPI/DNA staining is blue. Staining indicates that CAF colonies are either MMP1 positive or MMP1 negative. **F and G)** Two CAF lines were treated with LPS or TNF **(F)**, or with increasing concentrations of FBS **(G)**. The fraction of MMP1-positive cells was then determined by immunofluorescent staining.

To better understand MMP regulation in CAFs, we quantified MMP RNA expression in eight CAF lines derived from eight CRCs. The cultured CAFs expressed most of the MMPs observed in the single-cell RNA-seq analysis: MMP1, MMP2, MMP3, and MMP14 (Fig 2C; Fig 1C). Interestingly, MMP11 expression was not detected in the cultured CAFs, suggesting that this fibroblast subtype is under-represented in the cultures. Among the CAF-expressed MMPs, MMP1 showed the greatest variability. Spatial transcriptomic data from two independent CRCs showed MMP1-expressing CAFs to be clustered and variable in number, supporting MMP1 expression variability (Fig 2D). MMP1 mRNA levels in the CAF lines corresponded closely to MMP1 protein levels, indicating that MMP1 regulation is primarily transcriptional (Fig S3). Immunofluorescence staining of cultured CAFs likewise revealed an MMP1-expressing subpopulation (Fig S4). To determine whether MMP1 expression constituted a stable CAF phenotype, several CAF lines were plated at low density and allowed to proliferate to form discrete colonies. Immunofluorescent analysis showed that these colonies were either MMP1-positive or MMP1-negative. Fig 2E shows two examples where MMP1-positive and negative colonies were in close proximity. Given the size of the colonies, we estimate that the MMP1-phenoptype is stable through at least 5 cell divisions. The number of MMP1 staining cells varied between isolated CAF lines, ranging between ∼5 and 14% (Fig S5). To determine whether the number of MMP1-expressing cells changes following stimulation, cultures were treated with TNF, LPS or increasing levels of serum, followed by MMP1 expression analysis. TNF and high serum concentrations were able to increase the fraction of MMP1-expressing cells to a degree, whereas LPS did not (Figs 2F and 2G). These data show that stimuli can increase the number of MMP1-expressing CAFs, but that the number of cells that can be converted is limited. Overall, these data indicate that MMP1-expression is a relatively stable CAF phenotype that persists upon cell division and that this type of CAF varies between different CRCs.

### MMP expression in myeloid cells

Analysis of the myeloid-associated MMPs 9, 12 and 14 in the scRNA-seq data was performed focusing on three major myeloid cell types: monocytes, macrophages, and dendritic cells. The largest cluster (cluster 0) showed expression of all three myeloid-preferred MMPs in all three of the myeloid cell types (Fig 3A). Clusters 1, 2, 3, 5 and 6 showed relatively low MMP expression and were predominately populated with monocytes (Fig 3A). Two other clusters of interest are cluster 4, in which monocytes and macrophages express MMP14 almost exclusively, and cluster 7, in which macrophages and dendritic cells express high levels of MMP 12. To better understand the basis of MMP expression in the myeloid cells, high-expressing cells were resolved from low-expressing cells (Fig S6). Significant overlap in expression of the three myeloid MMPs was observed, consistent with an underlying coregulation (Fig 3B). Genes associated with high MMP 9, 12, or 14 expression were analyzed for enrichment using the Gene Ontology database (GO). All three of the MMPs were associated with the expression of genes involved in secretory vesicle formation and lysosome activity (Fig 3C). Based on this analysis, MMP expression in myeloid cells does not appear to be highly cell-type specific but is instead associated with a more robust expression of endomembrane system components, including secretory vesicles and lysosomes.

**Figure 3:**
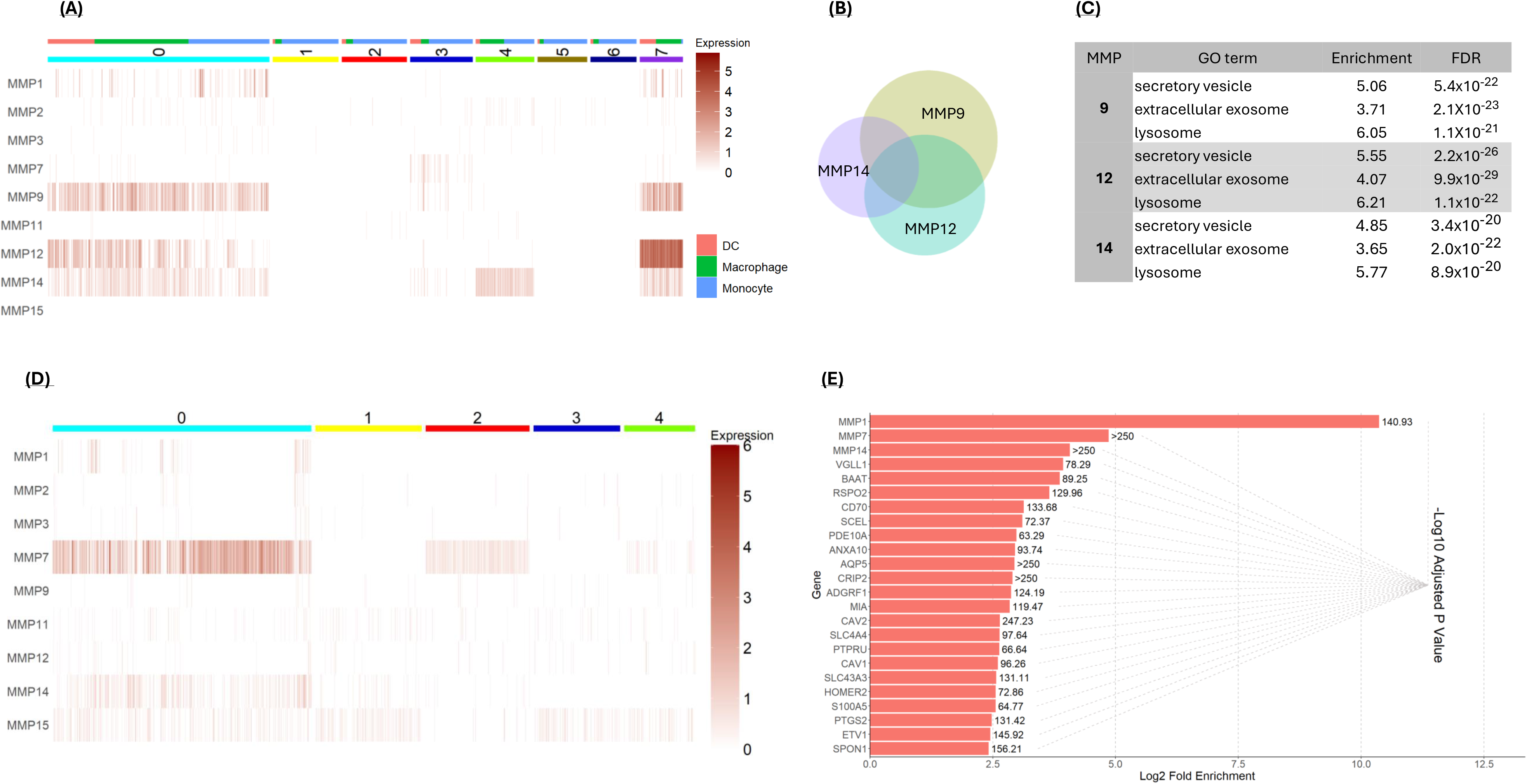
A) MMP-based clustering of myeloid cells from CRC. Monocytes, macrophages and dendritic cells (DC) from CRC were clustered based on their MMP expression to a resolution that generated 8 clusters. The cell type is indicated at the top of the graph. **B)** Overlapping expression of MMP 9, 12 and 14 in cancer-derived myeloid cells. The size of the circles and the area of overlap are proportional to the number of MMP-expressing cells. **C)** Myeloid cells expressing MMPs 9, 12 or 14 show significant enrichment for genes associated with secretory vesicles, extracellular exosomes, and lysosomes, based on the Gene Ontology (GO) Biological Processes database. **D)** Cancer cells were clustered based on MMP expression at a resolution that generated 5 clusters. MMP7 expression was most robust in cluster 0. **E)** The most enriched genes in cluster 0 versus all other clusters (based on fold-enrichment). The graph shows the log_2_-fold enrichment and the -log_10_ of the adjusted *p*-value).

### MMP expression in cancer cells

The scRNA-seq data showed that MMP expression is low in normal colonic epithelial cells but increases in colorectal cancer, with MMP7 showing high expression in a subset of cancer cells (Fig 1C)[10, 11]. As shown in Fig S7, MMP7 expression in cancer cells varies by several orders of magnitude. To better understand the nature of MMP7-expressing cancer cells, cancer cells were clustered based on their MMP expression. Cancer cells expressing the highest MMP7 were found in cluster 0 (Fig 3D). To determine how MMP7 expression correlated with other cancer cell characteristics, genes enriched in cluster 0 were identified (Fig 3E). The most highly enriched genes in this cluster were MMP1, MMP7, and MMP14. This finding indicates that these cancer cells likely possess high ECM-degrading potential, possibly affecting their malignancy [10, 11]. Additionally, this cluster shows enriched expression of the Wnt-signaling enhancers RSPO2, SPON1, and PTGS2 [12–14], and the Wnt target genes MMP17, PTGS2/COX2 and ETV1 [15–17]. This observation is consistent with reports that MMP7 is itself a Wnt target gene and suggests that this subpopulation of cancer cells has enhanced Wnt signaling [10, 18, 19].

### Spatial MMP expression in CRC

To determine how different MMPs might interact within a CRC, we analyzed spatial transcriptomic data obtained though 10X Genomics. The first cancer analyzed (tumor 1) was stage IIA CRC (sample ID CytAssist_11mm_FFPE_Human_Colorectal_Cancer). Moderate resolution data with approximately 20 cells per pixel was used to detect regions of MMP expression overlap. Analysis of spatial transcriptomic data showed distinct regional expression patterns for individual MMPs (Fig 4A). To understand how MMPs expression overlaps within the lesion, pixels were clustered based on MMP expression (Fig 4B) with these clusters “painted” on the tissue section (Fig 4C). The MMP expression clusters were driven largely by the cell types present in the pixel (Fig 4C). The MMP clusters also generated a patten similar to that made by the top differentially expressed genes (Fig 4C), a finding consistent with MMP expression helping to establish discrete subdomains. Cell marker analysis showed that most cancer cells were in MMP clusters 0 and 4 (EPCAM-expressing cells), whereas most fibroblasts were in cluster 1 (ACTA2 cells) and most inflammatory cells were in cluster 3 (CXCL8-expressing cells)(Fig 4D). Gene enrichment analysis of the MMP clusters generally confirmed these assignments (Fig 4E).

**Figure 4:**
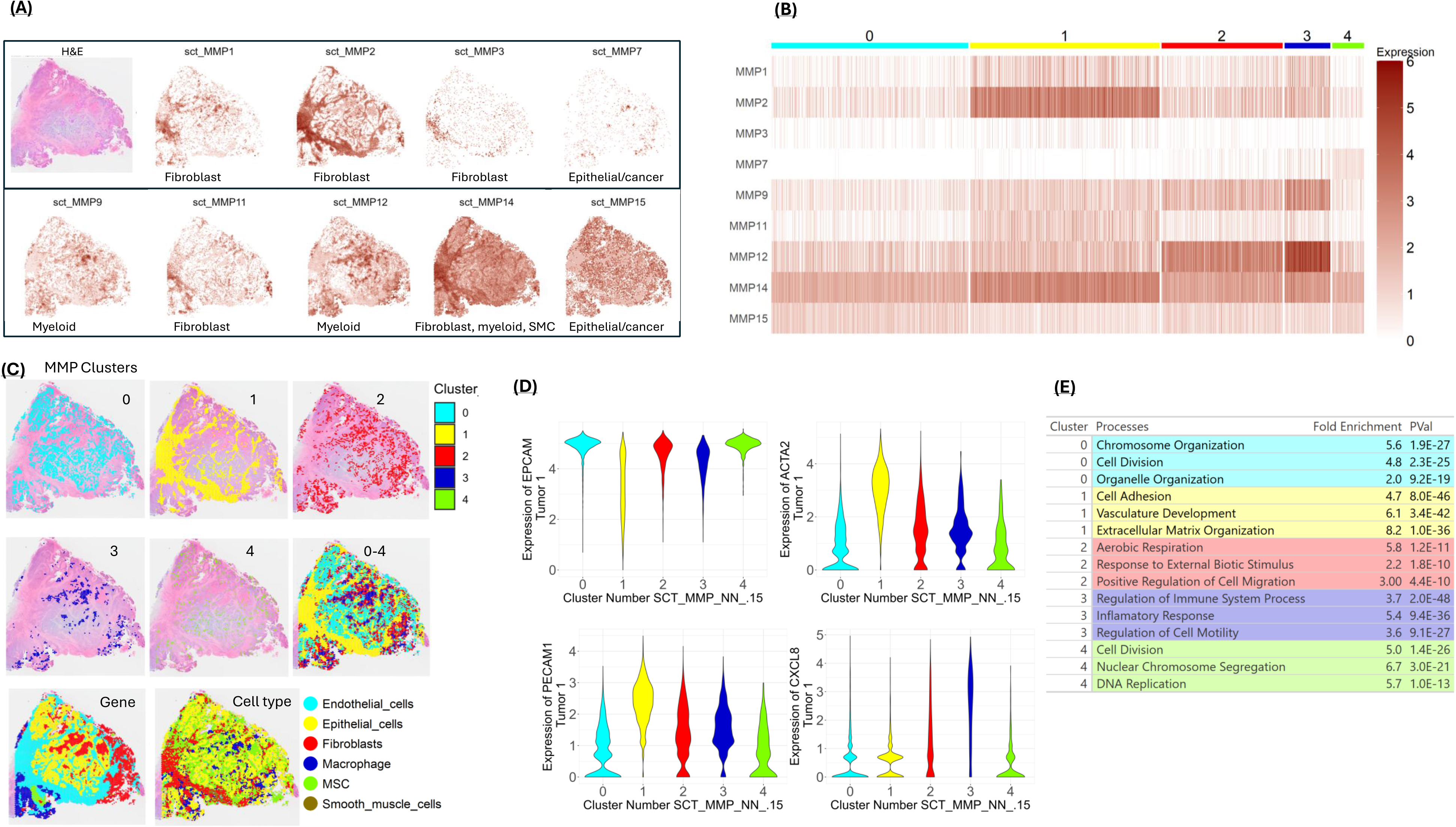
A) MMP expression mapped on a colon cancer tissue section referred to as “tumor 1”. An H&E-stained section is shown in the upper-left panel. The primary cell types associated with the different MMPs (from Figure 1) are indicated at the bottom of each panel. **B)** MMP expression patterns in the pixels of tumor 1 were clustered to generate 5 groups. **C)** MMP clusters from Figure 4B displayed on the tumor section, both individually and combined (0-4). Also shown are clusters based on the most differentially expressed genes (Gene-based) and Cell Type. MMP clusters resemble clusters generated from differentially expressed genes and cell type clustering. **D)** Violin plots showing selected marker genes in the MMP clusters. Epithelial/EPCAM-positive cells are a major component of clusters 0, 2, and 4, the ACTA2 activated fibroblast marker is highest in cluster 1, the angiogenesis marker PECAM1 is highest in cluster 1, and expression of the CXCL8 chemokine is highest in cluster 3. **E)** Upregulated marker genes for each cluster were passed through GO to determine upregulated paths and processes in each cluster.

A second CRC was analyzed for spatial MMP expression (tumor 2; obtained through 10X Genomics, sample ID Visium_HD_Human_Colon_Cancer). Although information on tumor stage was not available, examination of the tissue section revealed nests of neoplastic cells forming gland-like structures extending into the serosa, indicating a minimum progression level of stage II (not shown). The tumor 2 tissue section H&E showed extensive regions of smooth muscle and normal crypts, which were also revealed in the transcriptome analysis (Fig S8). To be consistent with tumor 1, these non-cancerous tissue regions (i.e., normal epithelium and muscle) were excluded from the analysis. Mapping of MMP expression on tumor 2 showed patterns similar to those found in tumor 1, with MMP expression dictated largely by the location of different cell types (Fig 5A). MMP clustering for tumor 2 is shown as a heat map in Fig 5B and “painted” on the tissue in Fig 5C. Cancer cells were found to predominate in clusters 0 and 1, with cluster 1 showing higher MMP7 expression and greater proliferative activity (Fig 5D)[20–22]. As in tumor 1, dense stroma formed a distinct MMP2-high cluster (cluster 2) colocalizing with high MMP11 and MMP15 expression, and the myeloid-expressed MMP9 and MMP12 grouped together in an immune/inflammatory cluster (cluster 3)(Fig 5D)

**Figure 5:**
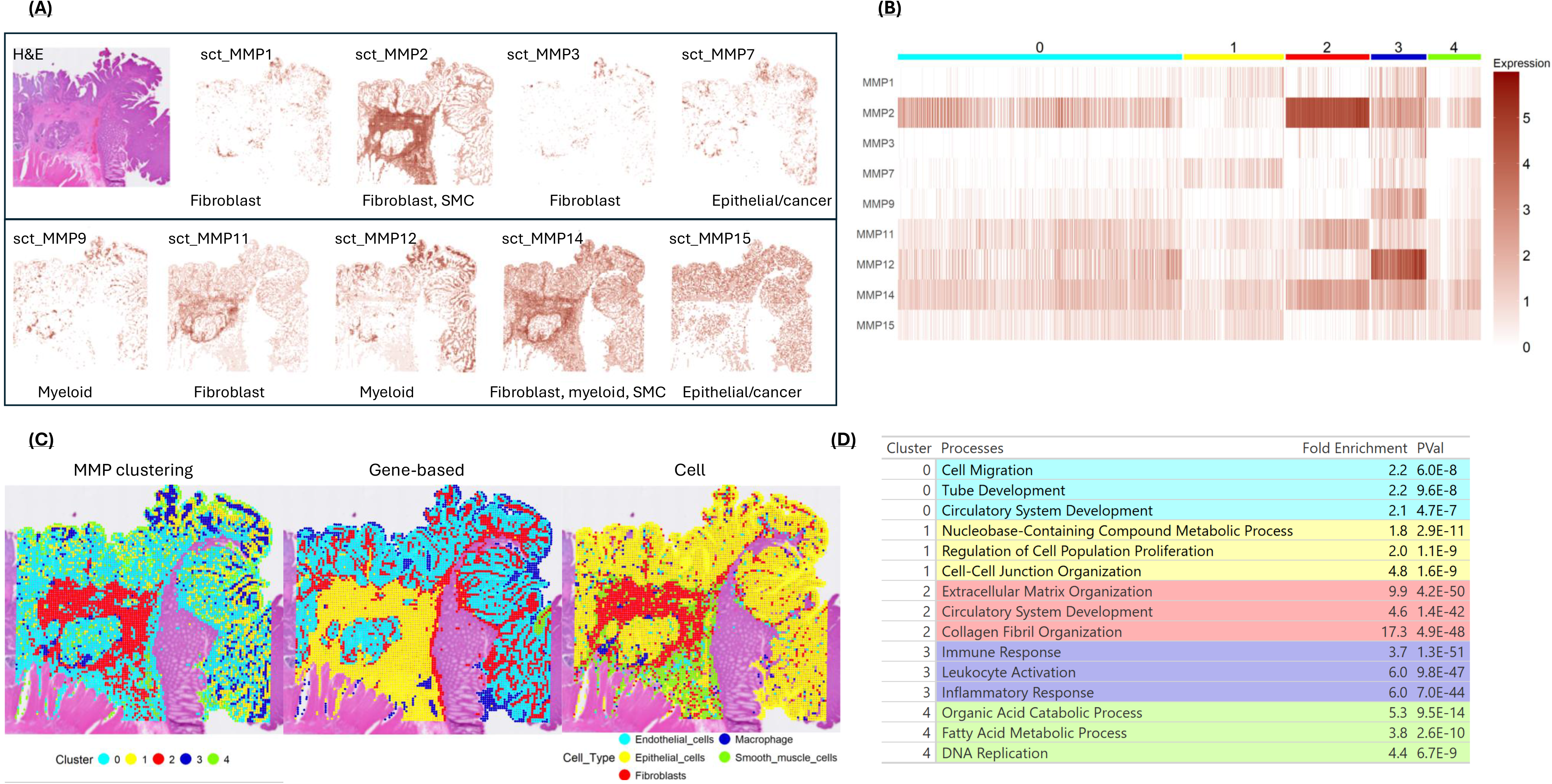
A) MMP expression mapped on a section of “tumor 2”. An H&E-stained section is shown in the upper-left panel. The primary cell types associated with the different MMPs (from Figure 1) are indicated at the bottom of each panel. **B)** MMP expression in tumor 2 clustered at a resolution to generate 5 clusters. **C)** MMP clusters from Figure 5B displayed on the tumor section. Also shown are clusters based on the most differentially expressed genes (Gene-based) and Cell Type. Note the MMP clusters show a similar pattern to the gene-based and cell type clustering. **D)** Differentially regulated genes from each MMP cluster were passed through GO to reveal the processes upregulated within the cluster.

Despite similarities with tumor 1, two differences were noted: 1) whereas the stromal region of tumor 1 showed high levels of MMP1 expression, relatively little MMP1 expression was found in the stromal region of tumor 2, and 2) tumor 2 showed a cluster with extensive MMP overlap, where fibroblast (MMP1, MMP3), myeloid (MMP9, MMP12, 14), and cancer cell (MMP7) enzymes colocalized (cluster 3) (Fig 5B). The overlap of MMP expression in cluster 3 suggests an area with high ECM remodeling potential, which may facilitate ECM dynamics and cell movement. This may be particularly relevant since MMP7-expressing cancer cells are also found in this cluster and may be more prone to move through the ECM.

### High resolution analysis of MMP7 expression

The spatial data above was analyzed at a moderate resolution (∼20 cells/pixel) to reveal subdomains of MMP-expression overlap within the tumor. However, tumor 2 data was available at high resolution (approximately 1 cell/pixel), allowing for a more granular, “single-cell” analysis. We focused on MMP7 since its expression by cancer cells is likely an important driver of cancer malignancy and metastasis [23–26]. The high-resolution spatial transcriptome data was filtered to identify pixels that include at least one MMP7 transcript. MMP7-expressing pixels localized near to the luminal surface as well as to stroma-encapsulated cancer cell “nests” deeper within the lesion, with differential gene expression grouping cells into distinct clusters, most notably with cluster 0 localizing to the luminal surface and cluster 1 encapsulating deeper tissue (Fig 6A). Analysis of MMP expression across these clusters revealed the expression of multiple MMPs in the stroma-encapsulated cluster 1 (Fig 6B). Gene enrichment analysis indicated that cluster 1 was enriched in genes involved in cell motility and ECM organization suggesting that this multi-MMP expressing cluster may define an invasive edge of the cancer (Fig 6E).

**Figure 6:**
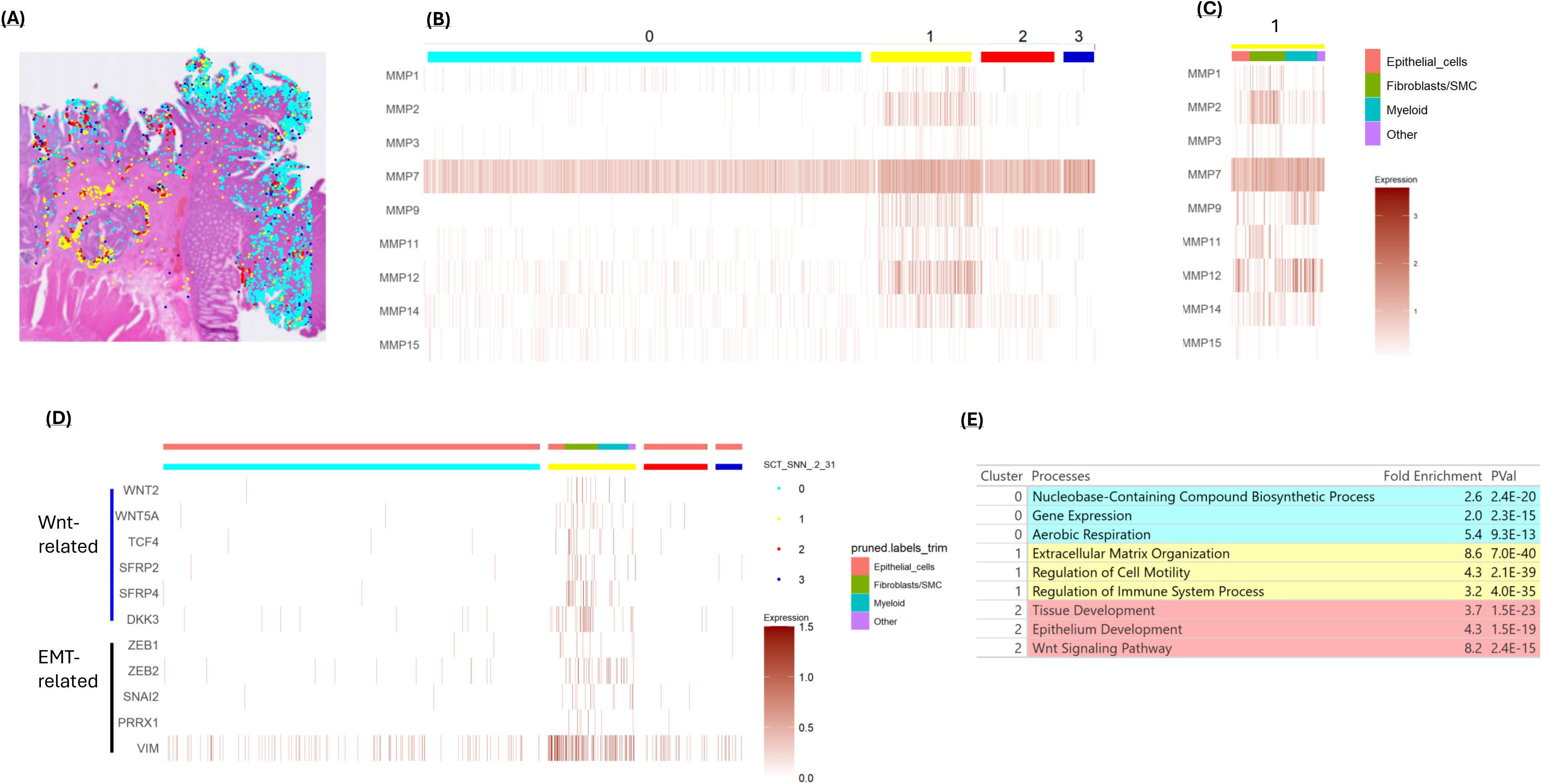
High resolution MMP7-expressing pixels were identified and sorted into 4 clusters based on differential gene expression. These pixels are displayed on the tumor section **(A)** and on a heat map **(B)**. Cells in cluster 1 tended to express multiple MMPs. **C)** Cell marker analysis indicates that cluster 1 is comprised of a number of different cell types. These cell types are labeled at the top on the heat map. **D)** The expression of Wnt and EMT related genes in the clustered, high resolution MMP7-positive cells. **E)** Marker genes defining each high-resolution cluster were passed to GO to determine the relevant upregulated pathways appearing to define each group. Cluster 3 was omitted due to a lack of significantly upregulated genes in relation to the other clusters.

Cell-type assignments for MMP7 expressing cells showed that clusters 0, 2 and 3 were largely cancer cells, consistent with MMP7 being expressed predominantly by cancer cells (Fig 1B and S9). However, cluster 1 was a mixture of cell types, including cancer cells, CAFs, and myeloid cells (Fig 6C). The cells in cluster 1 predominantly expressed the MMPs characteristic of their cell type, in addition to MMP7. These cells also tended to express an expanded repertoire of MMPs. This data raises the question: why do the different cell types in cluster 1 express MMP7, which is predominantly a cancer cell MMP? We investigated two possibilities. First, Wnt is a known MMP7 inducer so these cells may be in an area of elevated Wnt signaling [19, 27]. Consistent with this possibility, we found that genes related to Wnt signaling are enriched in cluster 1, including Wnt2, Wnt5A and TCF4 (Fig 6D). There are also a number of Wnt signal modulators enriched in cluster 1, including SFRP2, SFRP4 and DKK3. Wnt signaling in cluster 1 may therefore be instrumental in activating MMP7 in different cell types. Another possibility is that some cells in cluster 1 are MMP7-expressing cancer cells that have undergone an EMT. Consistent with this possibility, the expression of classical EMT promoters and markers are enriched in cluster 1, including ZEB1, ZEB2, SNAI2 (SLUG), PRRX1, and VIM (Fig 6D). In general, the cells in cluster 1 all appear to express an expanded repertoire of MMPs, suggesting strong activating signals and/or highly receptive cells (Fig 6C). Overall, we have identified an intriguing cluster of cells at the stroma-cancer border with broad MMP expression capabilities that may play a pivotal role in cancer progression.

## Discussion

Here, we provide a comprehensive analysis of the cell-type-specific expression of MMPs in CRC. Distinct MMP expression patterns were observed across different cell types, and although individual cells can express MMPs outside their primary profiles, obvious preferences were observed. Within each major cell category, additional layers of specificity were uncovered, related to cell subtype, activation status, or a combination of both. Although many of our observations have been previously reported for individual cell types, our analysis serves to integrate this information while also providing an analysis of spatial expression patterns in CRC.

CAFs are high MMP expressors in CRC and can be resolved into multiple MMP-based clusters, analysis of which suggests that different CAF subtypes express distinct MMP combinations. This observation aligns with previous findings that CRCs contain multiple CAF types, including resident fibroblasts, fibroblasts derived from infiltrating mesenchymal stem cells, and cancer cells that have undergone an epithelial-to-mesenchymal transition [9, 28, 29]. Precisely how the MMP-based clusters identified here relate to the CAF subtypes described by others will require a more detailed analysis. In contrast to CAFs, all myeloid cell types express the same set of MMPs: MMPs 9, 12 and 14, although monocytes tend to express lower levels than macrophages and dendritic cells. MMP expression levels in the myeloid cells correlate with the expression of genes related to secretory vesicle production, which is the route through which MMPs are secreted. In this regard, the MMP expression profile of myeloid cells appears to be related to a cell’s secretory activity rather than their type or subtype. The MMP most characteristically expressed in cancer cells is MMP7, a matrilysin with a broad spectrum of substrates. The expression level of MMP7 cancer cells ranges several orders of magnitude, with expression detected in approximately one-third of all cancer cells. Notably, MMP7 has previously been reported to be a target of the WNT signaling pathway, a commonly dysregulated system in CRC [30–32]. Concordant with this possibility, three WNT signaling enhancers, RSPO2, SPON1 and PTGS2/COX2, are also expressed in the MMP7-high cluster, along with the WNT-regulated genes MMP14, PTPRU, PTGS2/COX2 and ETV1 (Fig 3E). [12, 33–38]. The signals regulating this suite of genes may be a central determinant of metastatic potential.

Analysis of CRC spatial transcriptomic data showed MMP clusters consistent with the cell type expression patterns derived from the scRNA-seq analysis. MMPs expressed primarily by cancer cells localized to epithelial regions of the lesion, while fibroblast-derived MMPs were enriched in stromal areas. Myeloid cell-expressed MMPs clustered in regions characterized by immune and inflammatory gene expression. Computational clustering based on MMP expression yielded subregions within tumors that overlapped with regions defined by differential gene expression and cell type. This finding supports the idea that MMP activity contributes to the establishment and maintenance of tumor subdomains. Our findings show that fibroblast/stromal expressed MMPs are elevated in regions of active angiogenesis (Fig 4D and Fig 4E), suggesting that fibroblast-derived MMPs may play an important role in new vessel formation [39]. Consistent with this possibility, it has been found that MMP1, which is expressed primarily by CAFs, cleaves and activates the endothelial cell PAR-1 receptor to drive blood vessel growth and vascular remodeling [40]. Likewise, spatial enrichment of the myeloid MMPs may play a role in establishing immune and inflammatory cell hubs within the tumor [41].

Comparing the spatial MMP expression patterns of two tumors revealed similar clustering patterns, including stromal, inflammatory, and cancer cell clusters, with some notable differences. One difference was the number of MMP1-expressing CAFs within the tumor. Analysis of CAFs isolated from lesional tissue of patients likewise showed a broad range of MMP1 expression. Considering the scRNA-seq data and the clonal expression of MMP1 in cultured cancer fibroblasts, MMP1-expressing CAFs appear to be a fibroblast type that populates and potentially proliferates within tumors to different extents. Understanding how differences in fibroblastic/stromal MMP1 expression impacts tumor progression will require additional analysis.

Perhaps the most intriguing difference between the two tumors analyzed was the extent of MMP expression overlap. Since MMPs activate each other, and are active on different ECM substrates, overlapping MMP expression is likely relevant to lesion progression. Although only two tumors were spatially analyzed, different degrees of MMP expression overlap were observed. High resolution spatial analysis revealed clusters of cancer cells, myeloid cells, and fibroblasts at the cancer-stroma interface, all expressing an expanded repertoire of MMPs (Graphical Abstract). Within these multi-MMP expression clusters, elevated levels of WNT2, WNT5A, and TCF4 expression were also detected. Previous studies have reported that WNT signaling activates the expression of MMP7 and other MMPs, suggesting a potential mechanism for MMP activation within these clusters [30–32, 42–45]. Another feature of the multi-MMP cluster was the enrichment of the EMT markers ZEB1, ZEB2, and SNAI2 [46–48]. These zinc-finger transcription factors are instrumental in suppressing CDH1 expression, depolarizing epithelial cells and promoting their motility [49–51]. Similarly, the transcription factor PRRX1, which promotes the adoption of an activated fibroblast-like phenotype, was also expressed at high levels in this cluster [52]. The multi-MMP cluster appears poised for enhanced motility, in agreement with the GO Biological Process enrichment findings (Fig 6E). While analysis of additional tumors with accompanying clinical data would be required to fully understand the implications of MMP spatial overlap, our findings suggest that multi-MMP expression clusters driven by WNT signaling and active EMT may be of central importance to malignancy and metastasis.

There are several caveats to our study, the most obvious being that it is limited to RNA expression analysis. This limitation is particularly important for MMPs, as they can be regulated at multiple levels, including secretion, zymogen activation, and inhibition by TIMPs. Nevertheless, our findings have implications for understanding tumor progression and for the development of potential therapeutic strategies. Although MMP inhibitors showed promise in preclinical studies, most have failed in clinical trials because of excessive toxicity [53]. Broad-spectrum inhibitors such as batimastat and marimastat caused significant musculoskeletal toxicity, whereas more selective agents, such as DX-2400 (targeting MMP14), raised concerns because MMP14 is expressed in both tumor and normal tissues [54]. Understanding the spatial deployment of MMPs may aid in the development of selective inhibitors that minimize off-target effects. For instance, targeting MMPs expressed primarily in angiogenic regions (e.g., MMP1 and MMP3) may synergize with VEGF inhibitors to more effectively suppress neovascularization. Beyond the development of MMP inhibitors, MMP expression has also been proposed as a means of therapeutic targeting, either through localized drug release or antibody-directed approaches. MMP2-cleavable nanoparticles have been shown to successfully deliver drugs such as SN38 and doxorubicin in CRC models [55–58]. Our findings suggest that MMP7 could be exploited to deliver drugs directly adjacent to aggressive cancer cells [59]. More broadly, targeting the multi-MMP clusters identified in CRCs could provide a high degree of cancer specificity. Overall, there is still much to understand about the role, deployment, and regulation of MMPs in CRC. Such information could facilitate improved prognostic assessment and the development of novel therapeutic strategies.

## Data Availability

All datasets used in this study are publicly available at the repositories indicated in the Materials and Methods section.

## Acknowledgments

We thank Dr. Gaurav Joshi and Dr. Michael Thibodeau for their thoughtful comments and recommendations on this project.

## Ethics Statement

Colorectal tissue specimens used as a source of CAF cells were obtained from UConn Health/Waterbury Hospital. All samples were collected from consented patients undergoing colon resection and were de-identified in accordance with institutional protocols and established ethical guidelines.

## Abbreviations

Cancer associated fibroblast: CAF Colorectal cancer: CRC

Dulbecco’s Modified Eagle Medium: DMEM

Extracellular matrix: ECM Matrix metalloprotease: MMP Single cell RNA-seq: scRNA-seq Smooth muscle cell: SMC

Tissue inhibitor of metalloproteinase: TIMP

Tumor microenvironment: TME

Novelty and Impact: This study exploits recent advancements in spatial transcriptomics to explore matrix metalloprotease (MMP) expression distribution in colorectal cancers. MMP clustering is driven in part by cell type distribution within the lesion. Notably, regions of MMP expression overlap at the stromal border are observed, which may represent sites of cancer cell egress. This work provides insight into how MMPs shape consequential regions of the tumor microenvironment with prognostic and potentially therapeutic implications.

