## Supplemental Material for "Integrated Single-Cell and Spatial Profiling of Matrix Metalloprotease Gene Expression in Colorectal Cancer"

### Supplemental Figures

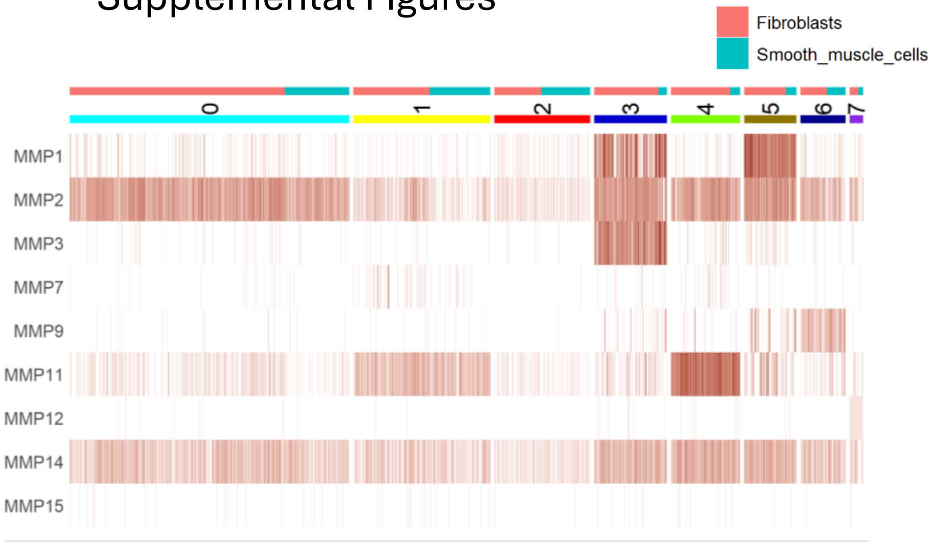


Figure S1: Cancer associated fibroblasts (CAFs) and smooth muscles cells from scRNA-seq data were clustered based on their MMP expression. Cells were also identified as fibroblast or sooth muscle based on cell using Single R.

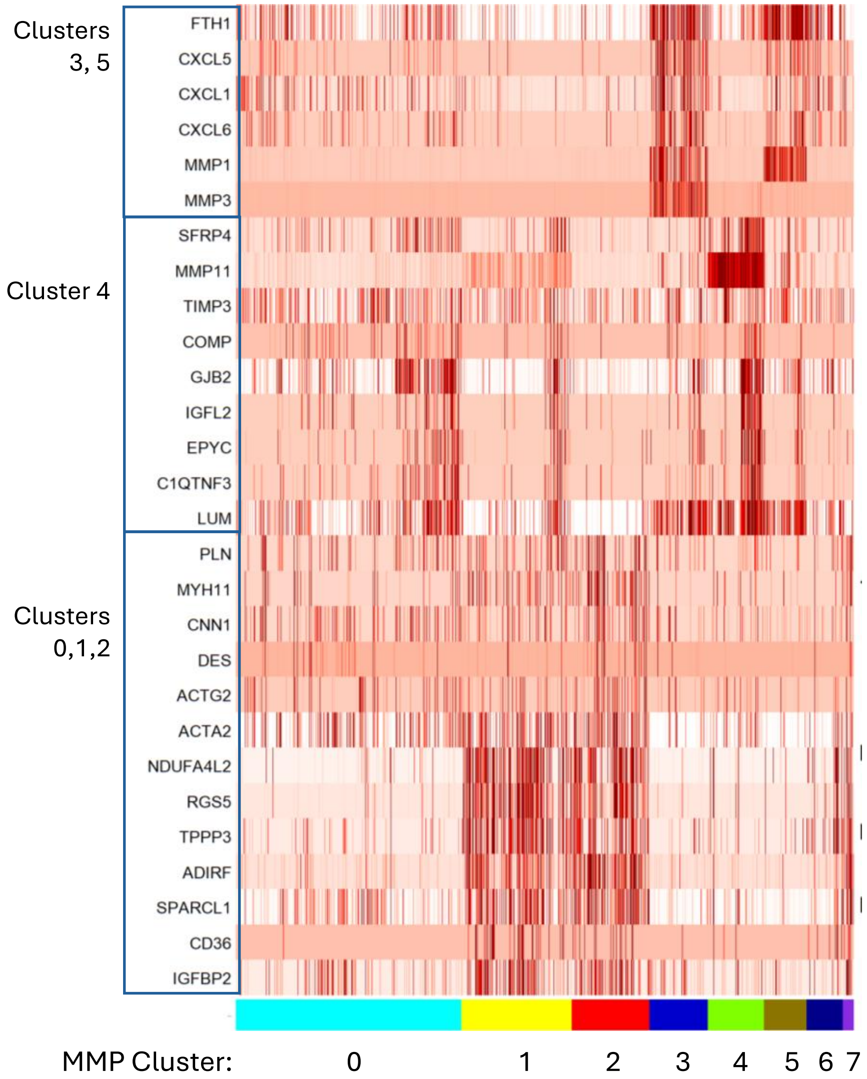


Figure S2: Differentially expressed genes in the MMP clusters observed in Figure S1 suggest different cell subtypes in the clusters.

### Supplemental Figures

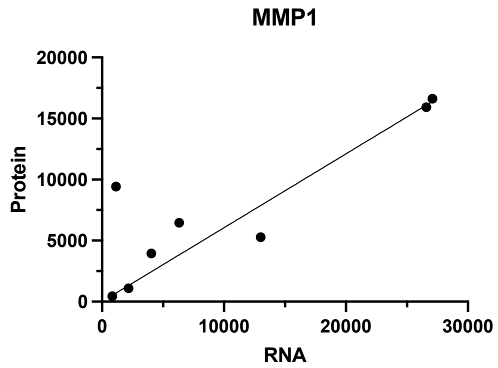


Figure S3: Relationship between MMP1 mRNA and protein expression in 8 CAF cell lines. RNA was quantified by RNA-seq and protein was determined by immunoblot analysis of secreted proteins.

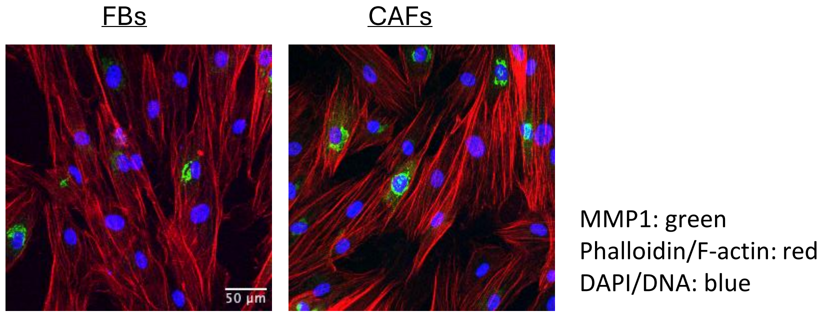


Figure S4: Fibroblasts from normal colon (FBs) or cancer (CAFs) were grown in culture and stained for MMP1, F-actin and DNA as indicated. images show a subpopulation of cells staining for MMP1 expression in both cultures.

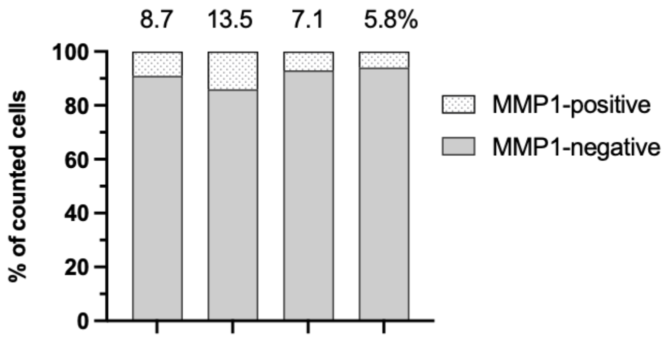


Figure S5: Percent MMP1-expressing cells in CRC cultured CAFs (from 4 independent CRCs).

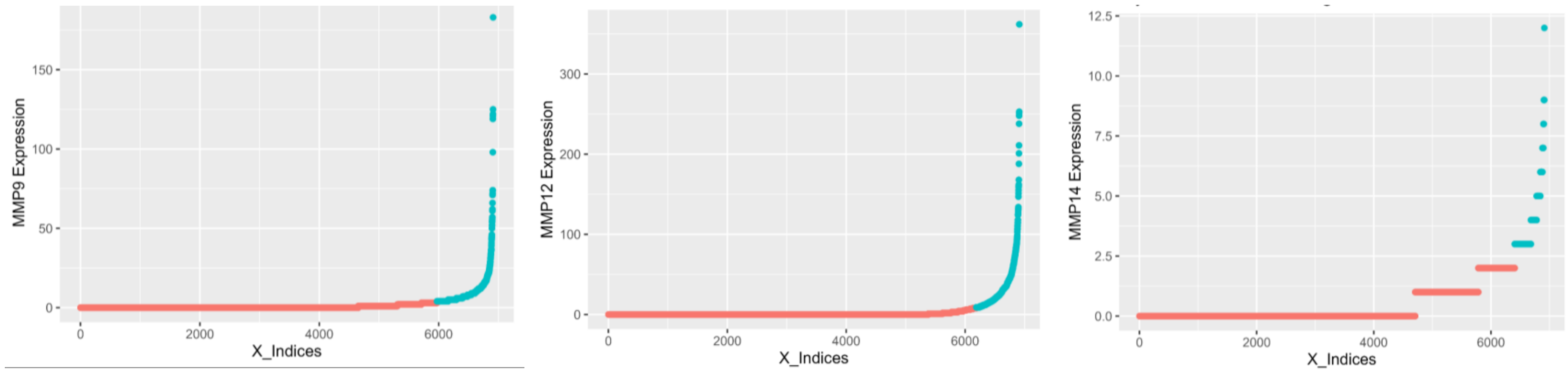


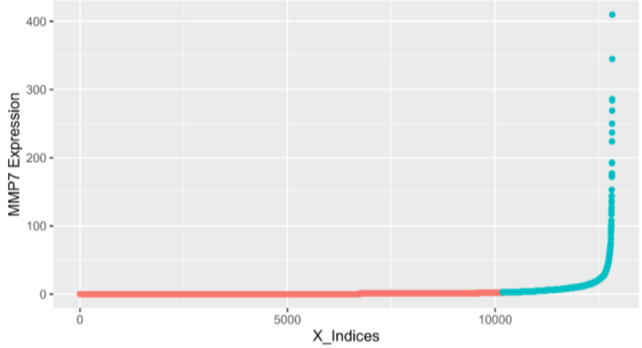


Figure S6: Expression MMPs in myeloid cells. Myeloid cells were pulled from CRC scRNAseq data. Cells positive for MMP 9, 12 and 14 expression were those cells with expression levels greater than the median. ”Positive” cells are shown in blue on the graphs.

Figure S7: Expression MMP7 in cancer cells. Cancer cells were pulled from CRC scRNAseq data. Cells positive for MMP7 expression were those cells with expression levels greater than the median. ”Positive” cells are shown in blue on the graph.

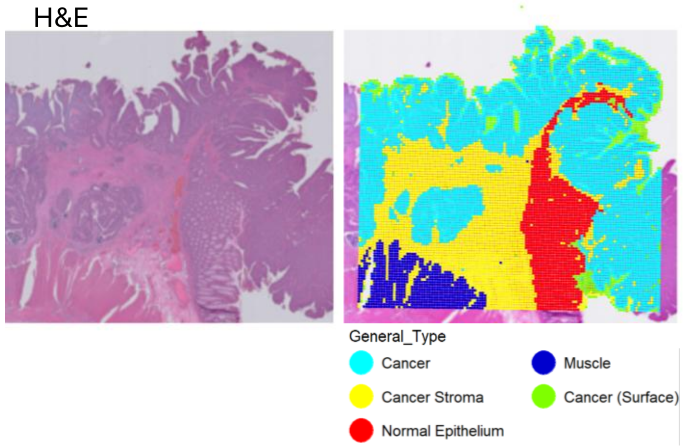


Figure S8: H&E staining shows cancerous tissue present in “tumor 2”. Non-cancerous regions were also revealed by gene expression analysis, which were removed prior to MMP cluster analysis for better comparison to tumor 1.

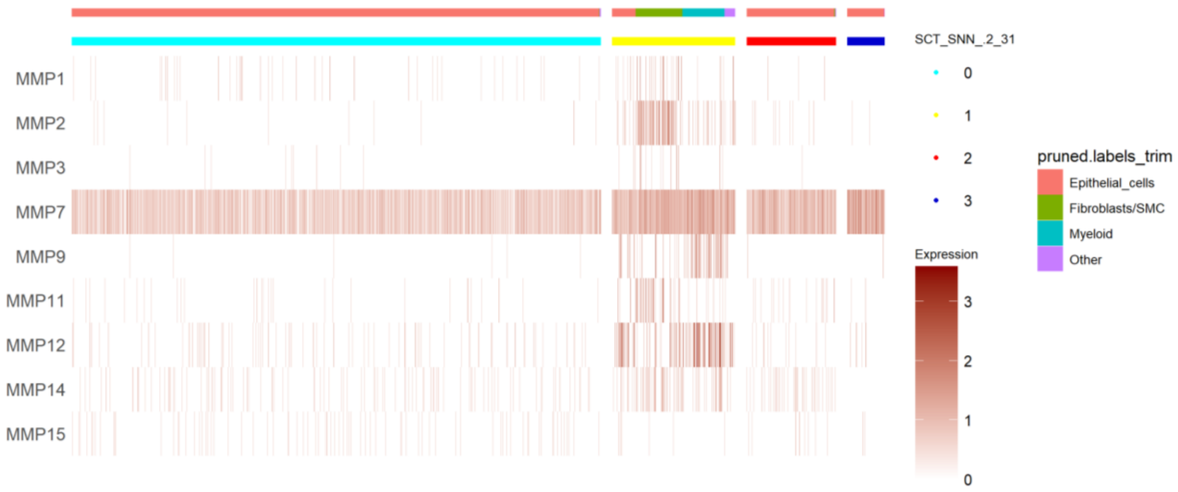


Figure S9: Full heat map for Fig 8C.
